# A note on the responsiveness of neurons

**DOI:** 10.64898/2026.10.21.760714

**Authors:** Hauke O. Wernecke, Arvind Kumar

## Abstract

The time constant of biological neurons is typically around 5-20ms. This raises the question of how a system made of slow devices can generate and respond to faster signals. A potential solution is a population of uncorrelated neurons that, as a whole, can respond arbitrarily fast despite individual slow neurons.

The response speed of a neuron or a population of neurons is often defined as how quickly neurons increase or decrease their firing rate given a change in the input, similar to a reaction time. Less emphasis is placed on the recovery time - the duration to transition and settle in a stationary state again in response to the change in the input.

Here, we investigate the reaction time and the recovery time of leaky integrate-and-fire neurons given a change in, either the mean input drive, or input fluctuation level.

In general, neurons react faster and fire quicker the larger the change in the input. Moreover, the closer the membrane potential is to the threshold, the faster the reaction. We observe that neurons react faster to increased input fluctuations than to stronger mean drive.

Complementary, the recovery time is systematically longer the larger the input change. Also, the closer the neurons’ potentials are to the threshold, the longer it takes the neuron to recover. Most interestingly, the baseline firing of the neuron strongly determines whether recovery is faster when the change happens in the mean drive or the fluctuation level. This observation suggests that communication via the mean drive and the fluctuation level could target different aspects of the overall response - quick reaction time and the recovery speed - based on the initial baseline activity.

## Introduction

The membrane time constant of biological neurons is considerably slow (5-20ms) (Destexhe, Rudolph, and Paré 2003). This raises the question of how a system made of slow devices can generate and respond to faster signals. A potential solution to this is that when neurons in a population are uncorrelated, the population as a whole can respond arbitrarily fast despite slow neurons (Ekeberg 1993; Tchumatchenko et al. 2011). In an *in vivo* setting, the balance of excitation and inhibition can render the population uncorrelated and thereby increase the response speed (Vreeswijk and Sompolinsky 1996).

When considering the response time of a neuron or population of neurons, the emphasis is often placed on how quickly neurons react to the change in the input, measured by the instantaneous increase (or decrease) in the output firing rate (Silberberg et al. 2004; Tchumatchenko et al. 2011).

Given a stimulus to a system or a neuron, the concept of the time scale can be interpreted as the time the system, or the neuron, takes until it reaches a steady state again. A neural population may exhibit a fast initial response, but the time to reach steady state may be slower – dominated by an ‘effective’ refractory period or even entail oscillatory activity (Knight, Omurtag, and Sirovich 2000; Ekeberg 1993; Tchumatchenko et al. 2011). Consequently, the population may display various activity responses to different perturbations or stimuli. Response transients pose a serious challenge for information representation in the neurons and can even cause large decoding errors (Lenninger et al. 2023).

Earlier investigation of reaction time assumed that the network was operating in a linear regime and neurons and synapses had a single time constant. This is far from reality. Even the simplest neuron models (e.g. leaky integrate-and-fire (LIF) neuron) display non-linearities, and hence can express various time scales depending on the operating point and the stimulus. On the far side of this scale, the dynamics of a biological neuron are shaped by a multitude of ion channels, and hence invariably display a temporal manifold (Marom 2010). Non-linearities may shape the temporal manifold and consequently affect the reaction time, as well as the recovery speed.

In this work, we focus on the recovery time, i.e., the time the population requires to settle in a steady state after a stimulus is presented. To this end, we injected Gaussian white noise (GWN) into neural populations and varied the noise parameters to mimic a stimulus. We found that recovery time was much longer than the reaction time. Interestingly, the non-linear reset mechanism is sufficient to introduce various time scales. The reaction time as well as the recovery time depended strongly on the baseline set point and the stimulus modality (whether the stimulus was a change in the input mean Δ*µ*, the variance Δ*σ* or both Δ*µ &* Δ*σ*).

When the stimulus pushed the neural population from baseline to the steady state, and then being relaxed again, the comparison of recovery times revealed that neurons exhibit a hysteresis in their activity. Furthermore, the presentation of transient stimuli exposed a non-monotonic relationship of the recovery time to both, the number of stimuli as well as the stimulus duration. These insights indicate the trade-off between fast, memoryless communication and history-dependent processing. Consequently, these findings suggest that the communication among neural networks may be finely tuned to typical activity (persistent or pulse-like stimuli) and connectivity characteristics (which affect the activity set point and the stimulus modality) of the transmitting population. Furthermore, the activity and connectivity can be adapted by neuromodulation in a context-dependent manner.

## Material & Methods

### Population of unconnected neurons

We simulated a population of 100 000 LIF neurons whose sub-threshold membrane potential *U* (*t*) dynamics were governed by

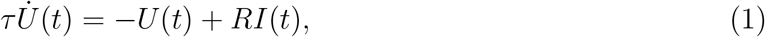

where *τ* = 15ms was the membrane time constant and *I*(*t*) was the external input current. The input resistance was *R* = 60MΩ. A neuron elicited a spike when the membrane potential crossed a threshold *θ* = 20mV, and then its membrane potential was reset to *V*_*r*_ = 0mV. After which, the neuron was refractory for t_ref_ = 2ms. The external current *I*(*t*) is the total input received by the neurons. All parameters of the network are given in Table 1.

**Table 1.**
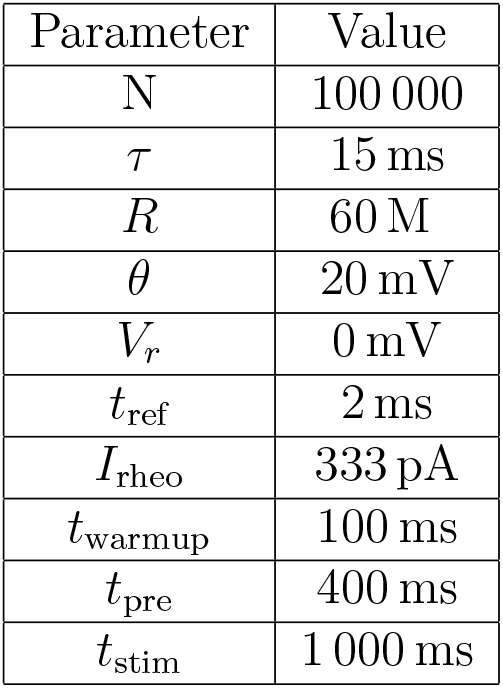
Neural parameter.

### Input

The external input *I*(*t*) consisted of a mean drive *µ* and the fluctuation level *σ*:

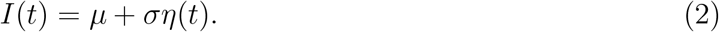

Here, *η*(*t*) was the Gaussian white noise with ⟨*η*(*t*)⟩ = 0 and ⟨*η*(*t*)*η*(*t*^*′*^)⟩ = *δ*(*t* −*t*^*′*^). Each neuron received an independent noise term. The input pushed the neural activity to the baseline firing rate (FR_pre_). At t_Δ_ = 500ms, mimicking the presence of a stimulus, the input was modified to reach the target activity FR_post_. The simulation continued for t_stim_ = 1000ms, which sufficed to reach a stable FR_post_.

A neuron can be driven to a firing rate by a pair of *µ* and *σ*, we refer to such a pair as set point. For the baseline activity FR_pre_, we set the mean drive *µ*_pre_ and derive the corresponding *σ*_pre_ utilizing the Siegert formula (Siegert 1951). Similarly, the fluctuation level can be set to obtain the required mean drive.

The stimulus altering the *I*(*t*) was either a change in the mean current Δ*µ*

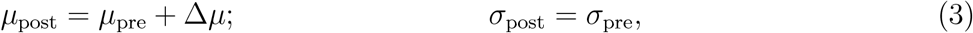

the fluctuation level Δ*σ*

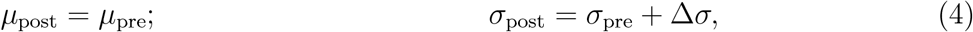

or both parameters Δ*µ &* Δ*σ*

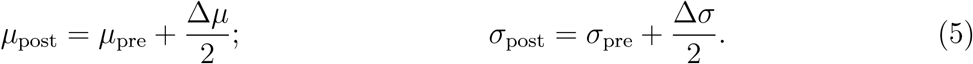

In case both parameters were changed, the magnitude of the change is half compared to individual parameter changes. We term these the three stimulus modalities.

### Reaction Time – Time to First Spike

The reaction time was quantified by the change in FR in response to a stimulus (Tchumatchenko et al. 2011). Specifically, the time to the first spike of each neuron was measured for t *>* t_Δ_.

### Recovery Time – Mean-Squared-Error-Reduction

The recovery time was determined as the time until the neural population reached steady state after presentation of the stimulus. Steady state was considered to be reached once transients were no longer detectable. In stochastic systems, such transient is called the initialization bias. A common approach to estimate the duration of the transient is the Mean-Squared-Error-Reduction (MSER) and its variations (White 1997; Franklin and White 2008). The idea of MSER is to discard initial *d* samples that are unrepresentative, as they are far from the sample mean. For a sequence of length *n*, the optimal truncation point *d*^*\**^ is found by minimizing the confidence interval (CI) around the sample mean:

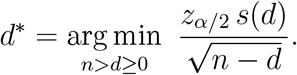

Here, *s*(*d*) is the large-sample standard deviation of the remaining sequence, and *z*_*α*/2_ is the value to calculate the 100(1 − *α*) CI. More explicitly, for a constant *z*_*α*/2_, the expression becomes

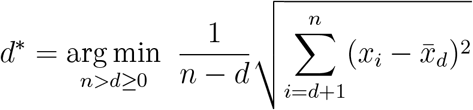

with 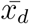 being the sample mean of the remaining sequence starting with sample *d* + 1. Note that the length of the whole sequence is large *n* ≫ 30, and the number of discarded samples is low, typically *d* ≈ 5% · *n*.

Moreover, the estimate of the confidence interval is biased for sequentially correlated observations which can be corrected. The application of the Philips-Perron correction, however, does not yield an improvement (Franklin and White 2008).

### Bootstrapping

In stochastic systems, bootstrapping is a technique to resample the observations to attain a distribution of estimates. However, the estimates are no longer uncorrelated since the same observation may be observed multiple times. Here, we split the neural population into subgroups of 500 neurons. Next, we pooled the spike times of neurons in *b* = 50 subgroups corresponding to a total of 25 000 neurons. Then, the peristimulus time histogram (PSTH; binwidth of 2ms) was derived, and the duration of the transient was estimated using MSER. We repeated this process *B* = 100 times to obtain individual trials. Importantly, the binwidth parameter had only a minor quantitative effect (Suppl. Fig. 1). When analyzing the recurrent neural networks, the excitatory populations were considered the subgroups. Here, the neural spike times of *b*_*RNN*_ = 10 subgroups were pooled, this is equivalent to a total of 50 000 neurons (*B*_*RNN*_ = *B*).

### Characterization of the Response Kernel

The neural activity was characterized as overshooting when the FR was higher than 1.1 FR_post_. Given the noise in the activity, many trials fulfilled this criterion. However, the distinction from an undershoot arises from the constraint that an overshooting kernel is required to be larger than the margin for an extended period, that is, for a minimum of 8 consecutive time points. Furthermore, the oscillatory kernel was based on at least one period that the activity is above the margin, as well as a following period of the activity being below 0.9 FR_post_.

### Recurrent Neural Network

The recurrent neural network (RNN) consisted of *N*_*E*_ = 5 000 excitatory and *N*_*I*_ = 1 250 inhibitory neurons, resembling a ratio of 4:1 (Braitenberg and Schüz 1998). The connection probability among the populations was *p* = 10% (Braitenberg and Schüz 1998).

In addition to the external input, the *i*th neuron of each population now also received recurrent synaptic current:

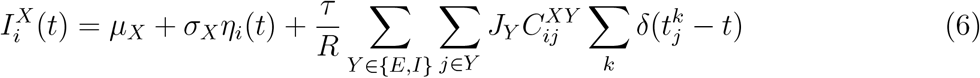

with *X* ∈ {*E, I*} marking the excitatory and inhibitory populations, respectively. 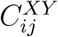 indicates a connection from the *j*th to the *i*th neuron. The time of the *k*th pre-synaptic spike is denoted by 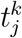.

A pre-synaptic excitatory spike evokes a jump in membrane potential of *J*_*E*_ = 0.025mV. The inhibitory synapses were *J*_*I*_ = − 8*J*_*E*_ stronger. Thus, the network operated in an inhibition-dominated regime. Moreover, the network exhibited asynchronous-irregular (AI) activity (Brunel 2000; Kumar et al. 2008). The FR of the inhibitory population was set to FR_*I*_ = 5Hz, driven by the external input with mean *µ*_*I*_ = 260pA (and corresponding fluctuation level *σ*_*I*_). The stimulus was only presented to the excitatory population to reach FR_post_. However, the change in the excitatory population changed the recurrent excitation. Therefore, the external input to the inhibitory population was tuned to maintain its FR, as well as its AI state. All parameters of the network are given in Table 2.

**Table 2.**
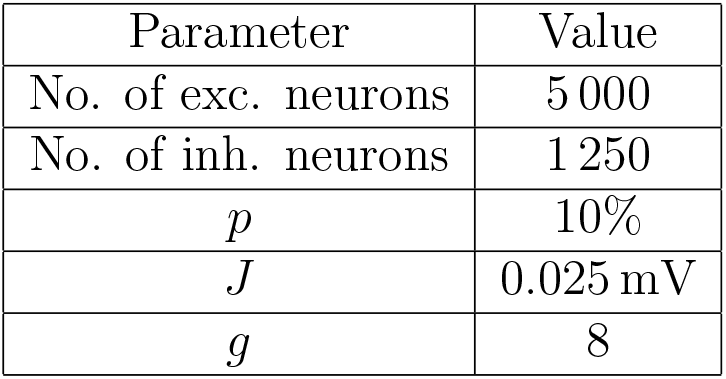
Parameter of the network and its neurons.

### Statistics

Statistical comparisons were performed by applying the one-sided Mann-Whitney U rank test (non-parametric) to evaluate whether the network activity recovered more slowly for the stimulus duration of t_stim_ = 15ms than the other stimulus durations. The significance threshold of *p <* 0.05 was adjusted using the Bonferroni correction for multiple comparisons (3 modalities, 3 numbers of stimulus pulses, 6 set points, 3 stimulus durations). The corrected significance thresholds were: 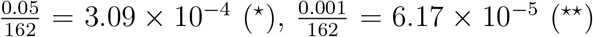, and 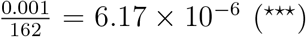. The recovery times for the stimulus duration t_stim_ = 15ms were compared to the other stimulus durations. The recovery was considered to be slower if the significance threshold was crossed in all three comparisons. Only the least significant *p* values are reported in the figure captions. No statistical method was used to predetermine the sample size.

### Simulation tools

All simulations were performed in NEST (Terhorst et al. 2025). The simulations and analyses were performed using custom python scripts.

## Results

Here, we are interested in characterizing how neurons as a population or as a part of a recurrent network respond to persistent or transient changes in the input. In particular, we want to quantify how much time neurons take to reach a steady state given a perturbation in their inputs.

Starting from time t_Δ_, we injected an external input into neurons already operating at a baseline firing rate FR_pre_ and measured when the neurons’ firing rates reached a stationary target activity FR_post_ (see Methods). We modeled the input to the neuron as GWN (Brunel 2000), and we considered three different types of input modalities:

1. Δ*µ*: the mean drive of the input is changed
2. Δ*σ*: the fluctuation level is adapted, or
3. Δ*µ &* Δ*σ*: both input parameters are altered.

### Response Kernel

First, we considered a scenario in which a single LIF neuron fired at baseline and received a persistent step-like stimulus (Fig. 1a). Irrespective of the stimulus modality (Δ*µ*, Δ*σ*, or Δ*µ &* Δ*σ*), we observed three different types of responses (Fig. 1b, c): (1) Undershoot: the neuron firing rate (FR) approaches FR_post_ asymptotically (Fig. 1c, left and middle column: red and yellow traces). (2) Overshoot: the activity briefly surges before decaying towards the steady state (Fig. 1c, left and middle column: turquoise traces). (3) Oscillatory Response: neural activity displays damped oscillations before reaching FR_post_ (Fig. 1c, right column: all traces). In our model, the shape of the response kernel depended on both the set point (the pair of *µ*_pre_ and *σ*_pre_) as well as the stimulus modality (Δ*µ*, Δ*σ*, or Δ*µ &* Δ*σ*). The response kernel shape varied smoothly as we changed inputs or the operating point. Based on these observations, we can argue that the neurons behave approximately like a second-order filter. Previously, all these response kernels have been observed in both experimental data and numerical simulations (Silberberg et al. 2004; Tchumatchenko et al. 2011; Khorsand and Chance 2008; Knight, Omurtag, and Sirovich 2000; Gerstner 2000).

**Figure 1.**
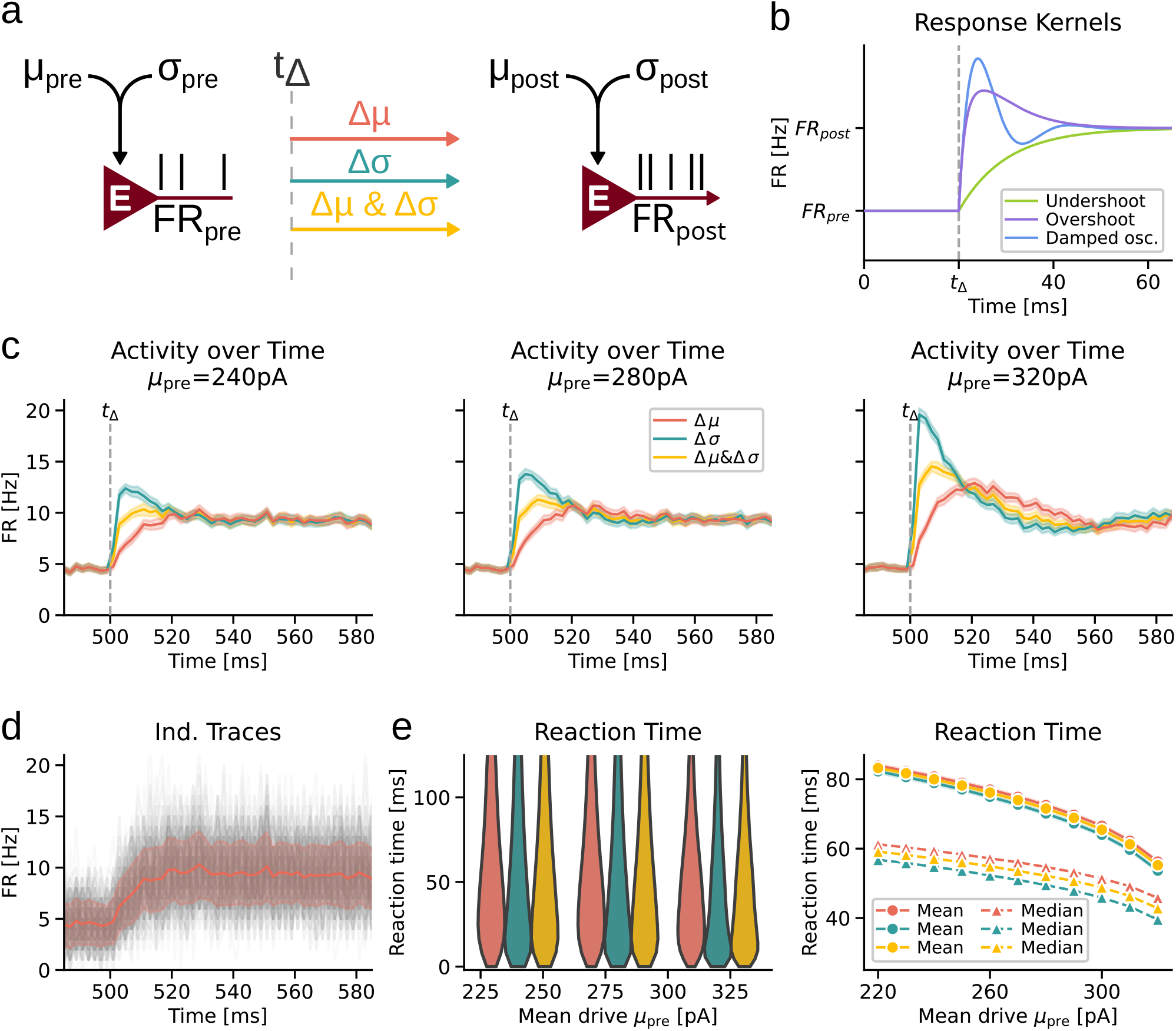
Network architecture, response kernels and reaction time **(a)** Network architecture: Unconnected neurons receive external GWN input. At time *t*_Δ_, the input is changed and the neurons are driven towards a target firing rate FR_post_. **(b)** Response kernels: three response kernels are observed across different setpoints (*µ*_pre_, *σ*_pre_) and the stimulus modality (Δ*µ*, Δ*σ* or Δ*µ &* Δ*σ*). **(c)** FR traces for three setpoints, increasing mean drive *µ*_pre_ from left to right. **(d)** Individual traces neuronal subgroups (*N* = 500 neurons) displaying large variability. **(e)** Reaction time: (left) distribution of the first spike per neuron after the stimulus was presented. (right) Mean and median of the distributions, respectively.

### Reaction Time – Time to First Spike

One part of the responsiveness of neurons is the time required until a neuron spikes after a stimulus is presented - we call this the reaction time. Here, we observed faster reaction times whenever the fluctuation level was increased as compared to the change in the mean drive of the stimulus (Fig. 1e). Not only was the average reaction time lower, but the distributions were more skewed towards faster reaction times when the fluctuation level was changed.

The baseline set point (the pair of *µ*_pre_ and *σ*_pre_) to the neurons also affected the reaction time. Increasing the mean input (while reducing the fluctuations to maintain the baseline firing rate) resulted in faster reaction time (Fig. 1e).

These observations are consistent with previous results that suggest fast signal transmission when the information is encoded in the input variance (Silberberg et al. 2004).

### Recovery time – Time to Stationary State

The second part of the responsiveness of neurons is the time to reach a steady state again - we call this the recovery time. Assuming that information is encoded in the FR of neurons, the responses must reach a steady state quickly for fast and accurate encoding of information otherwise transient fluctuations can impair fast inference of inputs (Lenninger et al. 2023).

In our model, the recovery time depended on the input modality (Δ*µ*, Δ*σ*, or Δ*µ &* Δ*σ*) as well as the difference between baseline and target FR (Fig. 2a, c). Interestingly, the variance of recovery times was larger when the fluctuation level was increased (Δ*σ*; see turquoise distributions in Fig. 2b). This phenomenon may arise from the generally stronger initial response (i.e., overshooting response kernel) for Δ*σ* (see Fig. 2a, c). Note that the variance of both baseline and target firing rates was similar across the input modalities during steady state (Suppl. Fig. 1a-c).

**Figure 2.**
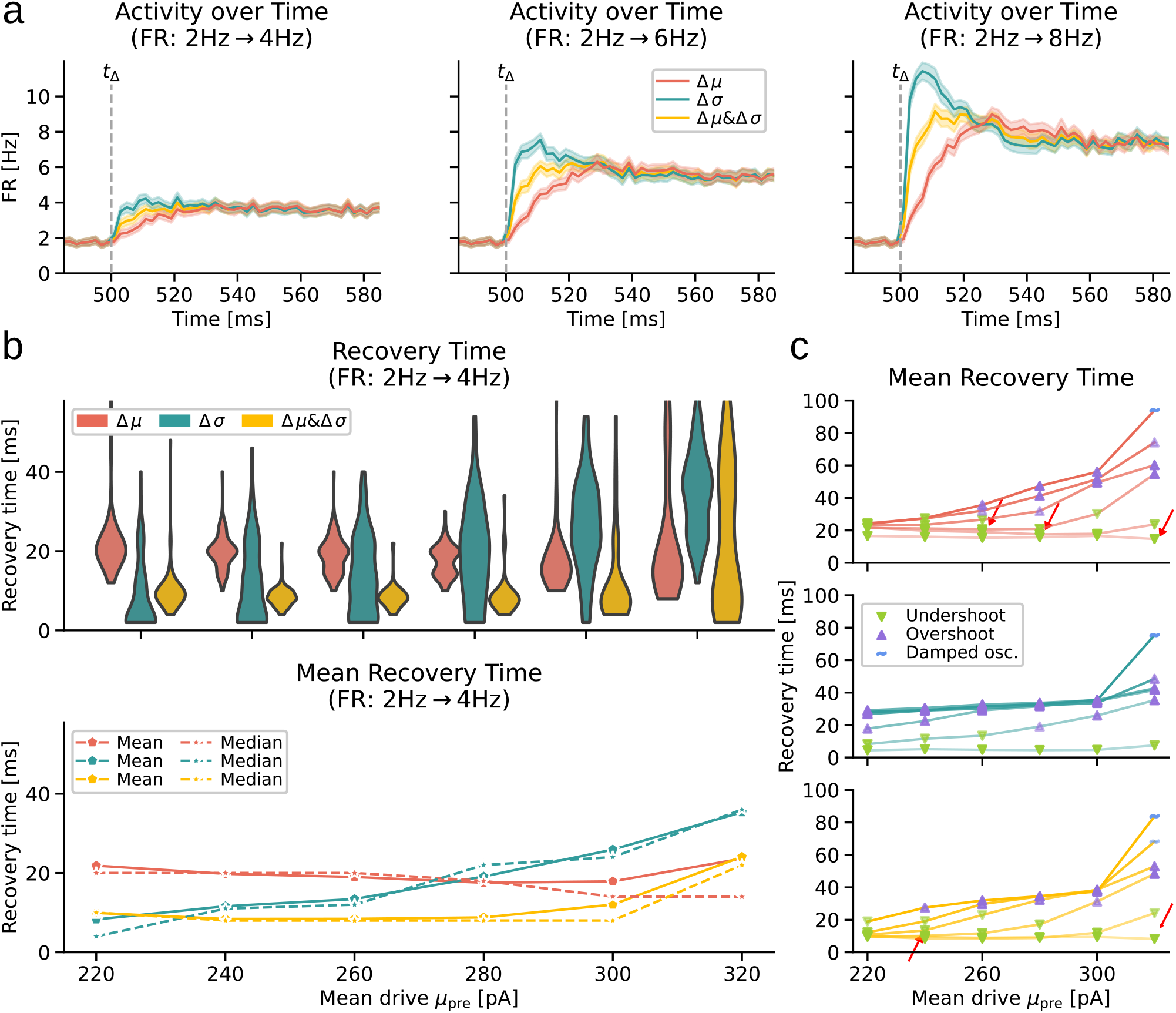
Recovery time across various set points. **(a)** Effect of the target FR_post_: Larger differences between the baseline and the target FR exhibit longer recovery times. Moreover, the response kernels changed qualitatively. From left to right: Increase in the target FR_post_. **(b)** Recovery time estimates: (top) Distribution of recovery times across various set points (*µ*_pre_ ∈ (220pA, 320pA)). (bottom) Mean (solid) and median (dashed) recovery times. **(c)** Mean recovery times across target FR_post_ (low alpha is FR_post_ = 3Hz, and high alpha is FR_post_ = 8Hz). Markers indicate the predominant response kernel(▼: undershoot/overdamped; ▲: overshoot/underdamped; : ∼ oscillatory response), and the stronger the alpha, the more prevalent the kernel. Red arrows indicate the mean drive *µ*_pre_ that results in the fastest recovery.

When the neurons were in a high-mean set point (i.e., high *µ*_pre_ paired with low *σ*_pre_), the recovery times were usually longer (Fig. 2b, c). This happened when the response kernel was of overshooting or oscillatory type - or transitioning towards it (Fig. 2c: response kernels indicated as markers, and their prevalence by the respective alpha value).

However, when the responses were undershooting (being overdamped), the minimum recovery time was observed for intermediate values of the mean drive (Fig. 2c: red arrows). While very weak *µ*_pre_ delayed the recovery, strong *µ*_pre_ shifted towards the overshooting (underdamped) response. Hence, a sweet spot was comparable to a critically damped second-order low-pass filter.

In general, large differences in the estimate of recovery time were seen when the response kernel changed qualitatively, e.g., from the asymptotic undershoot to a quick overshoot, or from the overshoot to an oscillatory pattern.

Lastly, towards high-mean (and low fluctuation) set points, the response kernels were predominantly of an oscillatory nature. Therefore, the recovery time distributions tended to become bimodal (Fig. 2b) as the steady state is identified either before or after the last low-amplitude cycle (Franklin and White 2008).

### Response to Pulse Trains

Naturally, step-like inputs are rare. Continuously varying stimuli or a set of brief discrete events are biologically more plausible. Here, we investigated how pulse trains consisting of one, two, or five discrete stimuli affected the recovery time of neurons. While a single stimulus may provide a metric for a neuron’s bandwidth, multiple stimuli would also emphasize history-dependent activity.

Again, neurons were driven towards a target FR_post_. However, after a stimulation period T_stim_, the input was reset to baseline. The edge case is a long stimulation period T_stim_, which is similar to the standard onset (5 →10Hz) and offset (10 →5Hz) dynamics, respectively (Fig. 3a). Input modality affected the onset and offset recovery times. For Δ*µ*, the offset recovery was faster (Fig. 3a: red ✦). For Δ*σ*, in contrast, the onset recovered more quickly (turquoise •).

**Figure 3.**
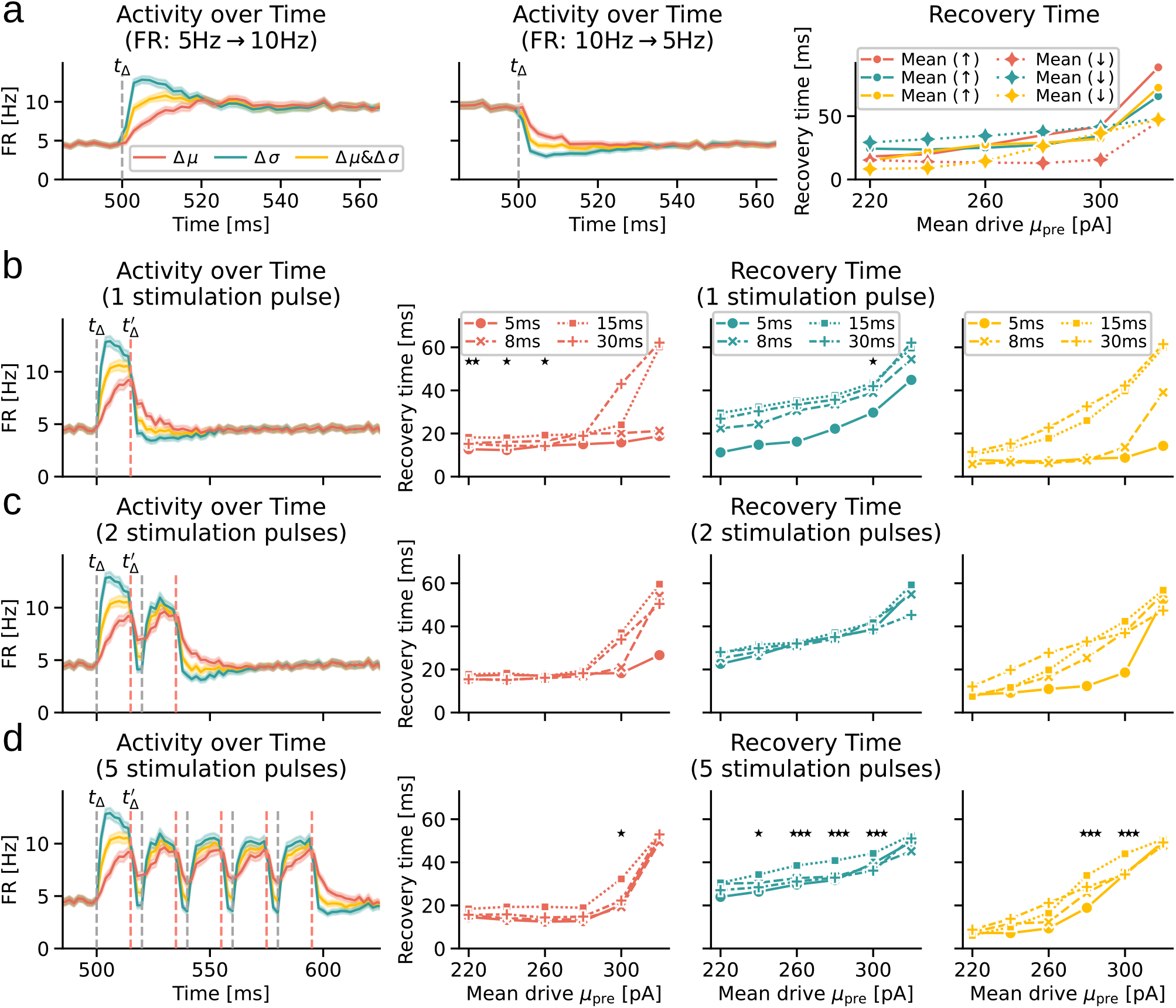
Recovery time for transient stimuli. **(a)** Onset (left) and offset (middle) of a persistent stimulus, and their respective recovery times (right) across all modalities; •: Onset, ✦: Offset. **(b)** One stimulus pulse: neural activity over time in response to a single stimulus pulse(left). Recovery times for a stimulus of varying duration (Δ*µ*: red; Δ*σ*: turquoise; Δ*µ &* Δ*σ*: yellow). **(c)** Same as (b) for two consecutive stimuli separated by T_pause_ = 5ms. **(d)** Same as in (b) for five consecutive stimuli.

Building on these observations, we reduced the stimulation period to shorter periods than the recovery time of the persistent onset stimulation (except T_stim_ = 30ms) and estimated the recovery time from the last offset of the pulse train.

A pulse-like stimulus revealed how stimulus duration also affected the recovery time had a non-linear relationship with the stimulus duration (Fig 3b). Next, we presented stimuli with two or five pulses (Fig 3 c, d). These experiments revealed that the recovery time also depended on the number of pulses, especially when the neurons were operating in a high-mean set point (high *µ*_pre_ and low *σ*_pre_). Although the underlying mechanism is unclear up to now, it was apparent that in several cases the recovery time varied non-monotonically with the number of pulses (Suppl. Fig. 2).

Furthermore, in a few cases, the recovery was the slowest for an intermediate pulse duration. This phenomenon was more noticeable when input was modeled as Δ*σ* or Δ*µ &* Δ*σ*, e.g., compare the recovery time for *T*_stim_ = 15ms (dotted; ■) and *T*_*stim*_ = 30ms (dash-dotted; +). Finally, the difference in the onset and offset dynamics and dependence of recovery time on pulse duration and count suggest that even unconnected neurons can and do exhibit hysteresis (Fig. 3; Deger et al. (2010)).

### Recovery Time of a Biological Recurrent Neural Network

Thus far, we focused on unconnected neurons. Next, we estimated the recovery time when neurons were part of a recurrent network of excitatory and inhibitory neurons (Fig. 4a; see Methods). We tuned the network (balance of excitation and inhibition) to operate in an asynchronous-irregular state (Brunel 2000; Kumar et al. 2008). Here, the stimulus was only presented to the excitatory population to drive the activity to FR_post_. Even the stimulated network’s activity remained asynchronous-irregular with a small increase in the Fano factor (weak correlations) for stronger external mean drive *µ*_pre_ (FF; Fig. 4e). At high mean set points (*µ*_pre_ ≥ 300pA), the network dynamics crossed a bifurcation point and displayed synchronous-irregular activity.

**Figure 4.**
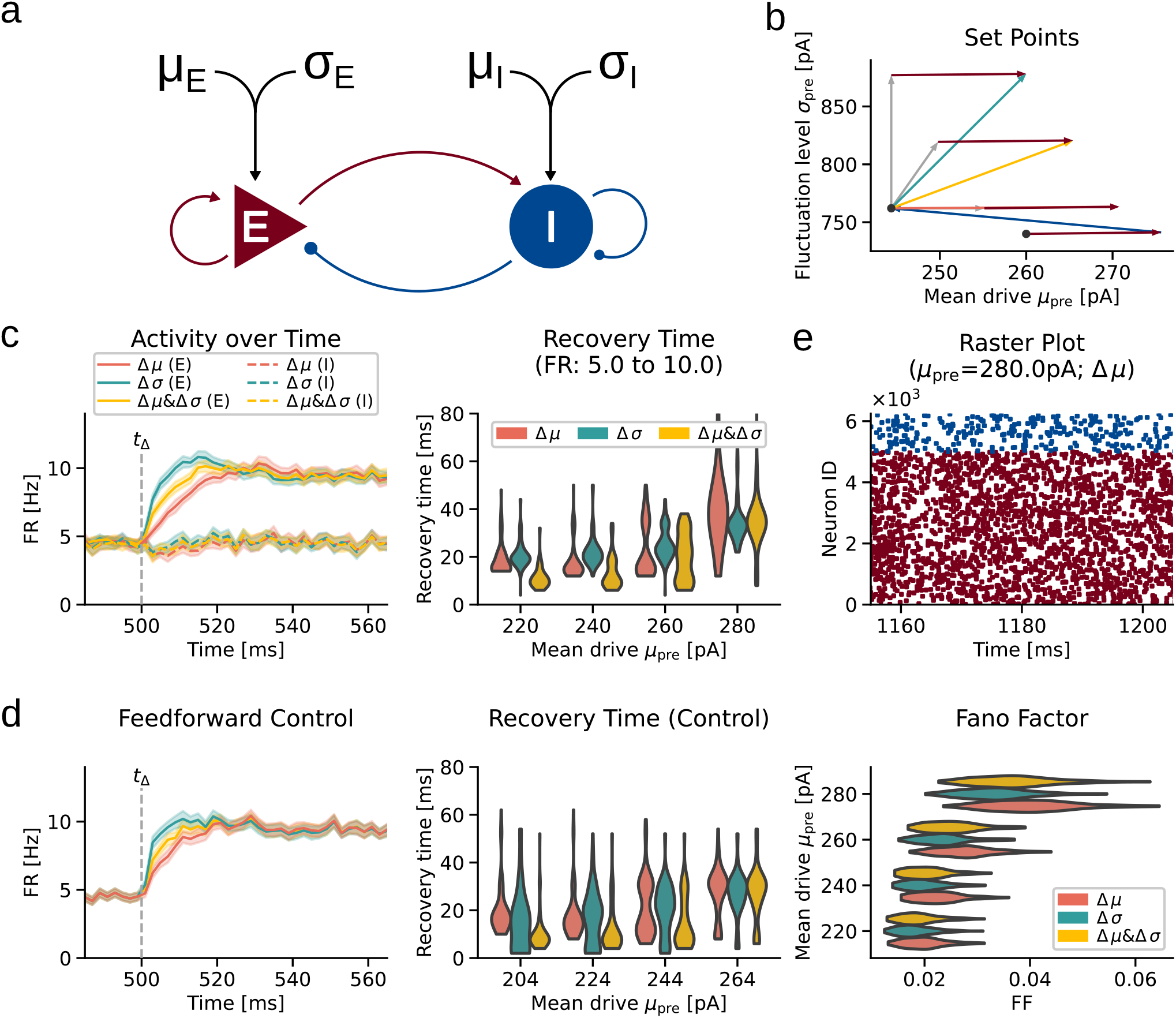
Recovery time in a recurrent neural network **(a)** Network architecture: The excitatory and inhibitory populations receive external input. Only the input to the excitatory population was stimulated (see Methods). **(b) (c)** (left) Activity of the excitatory (solid lines) and the inhibitory (dashed lines) populations (*µ*_pre_ = 260pA). (right) Distributions of recovery times across set points and stimulus modalities. **(d)** Same as in (c) for the unconnected population receiving feedforward external input. **(e)** (top) The network exhibited asynchronous-irregular activity in steady state. (bottom) The Fano factor was very low, and displayed a slight increase for higher *µ*_pre_.

Qualitatively, the same three response kernels were observed in the network responses (Fig. 4c left). Similar to the unconnected population, the recovery time strongly depended on the set point and the stimulus modality (Fig. 4c right).

While the response kernels were similar, there were two crucial differences in the network and unconnected neuron population. First, the recurrent network dynamics reduced the total mean drive (Sadeh, Cardanobile, and Rotter 2014). Consequently, the fluctuation level to attain the same FR was higher in the network than in the unconnected population. Hence, a recurrent network with the same external input operated at a set point with higher fluctuations than a comparable population of unconnected neurons (Fig. 4b).

Second, in the network setting, the neurons received two different inputs: (1) the external feedforward drive (*µ*_pre/post_ and *σ*_pre/post_), and (2) the time-varying recurrent synaptic input. To better understand the contribution of the time-varying input, we performed a control experiment: we calculated the total input current *I*_*SS*_ (into the exc. population) at steady state (FR_pre_ and FR_post_) and then injected *I*_*SS*_ into an unconnected population. Through this approach, we removed the time dependency (and the weak correlations).

We found that for matched response kernels, the recovery times were longer for the network than for the unconnected population. This is because of the time-varying component of the input received by the neurons in the network. When the response undershot, the current *I*(*t*) during the transient phase was lower than *I*_*SS*_. Therefore, the unconnected population reacted and recovered faster. On the other hand, an overshooting kernel caused temporarily stronger recurrent excitation (*I*(*t*) *> I*_*SS*_). This in turn led to a stronger overshoot and consequently, longer recovery times of the network activity compared to those of the unconnected population. In rare cases, the temporarily weaker recurrent excitation evoked an undershooting kernel compared to a (slightly) overshooting kernel in the control experiment. Then, the RNN exhibited faster recoveries.

These observations warrant cautious interpretation as weak correlation in the RNN may partly account for differences in recovery times. Moreover, the pooling of neurons differs for the RNNs and the unconnected populations. The resampling of larger subgroups (RNNs) causes the samples to be less independent than with small subgroups (control; see Methods).

## Discussion

The response properties of a neural population are commonly analyzed in terms of how fast neurons react, how strong, or even how accurate the response is. However, the time the neural population requires to reach a stationary state is rarely investigated. Here, we characterized the recovery time of leaky integrate-and-fire (LIF) neurons to persistent and transient stimuli.

First, we found consistently shorter reaction times when the stimulus was a change in the variance of the input current (Δ*σ* compared to Δ*µ*; cf. Silberberg et al. (2004) Next, we confirmed the presence of response kernels to persistent stimuli: being undershooting, overshooting, and oscillatory responses (Fig. 1; Silberberg et al. 2004; Tchumatchenko et al. 2011; Khorsand and Chance 2008; Knight, Omurtag, and Sirovich 2000). For a given stimulus modality, oscillatory responses were slowest to recover, while undershooting responses were the fastest. Quantitatively, the recovery time strongly depended on the set point (the pair of *µ*_pre_ and *σ*_pre_) and the stimulus modality (Δ*µ*, Δ*σ*, or Δ*µ &* Δ*σ*).

In general terms, the higher the *µ*_pre_, the more likely an overshooting or even an oscillatory response was. Interestingly, when the predominant kernel transitioned from an undershooting to an overshooting response (for increasing *µ*_pre_), we observed the minimum recovery times (Fig. 2c).

As neural populations are usually embedded into neural networks, we also studied recurrently connected networks. In our networks, the inhibition dominated the excitation, and thus displayed larger fluctuations with lower mean drive (Fig. 4; Sadeh, Cardanobile, and Rotter 2014). As for an unconnected population, we found that the recovery time of the network depended again on the set point and stimulus modality.

A crucial difference was that the total current input (feedforward and recurrent) in the network was activity- and thus time-dependent compared to the external input received by an unconnected population. For matched conditions (set point and modality), the network recovered more slowly than an equivalent unconnected population. This is due to the recurrent excitation in the network: the total input was lower than in steady state, resulting in a slower rise. When the activity is above the target FR_post_, it led to temporary over-excitation that supported a stronger overshoot.

Regardless of the nature of the stimulus, Vreeswijk and Sompolinsky (1996) proposed linear network responses based on large networks and cancellations of excitation by quick inhibition. Their results indicate low-pass filtered responses with a single time constant, and that implies a quick recovery. This neglects the second-order dynamics that LIF neurons display (Hahn et al. 2014). Moreover, their network operated in a (biologically plausible) fluctuation-driven regime. While we observed similar dynamics (especially for Δ*µ* and low *µ*_pre_), this might not be the optimal set point. The optimal set point resembles a critically damped second-order filter that balances quick reaction time and fast recovery (Fig. 2c). As we observed that RNNs often tend to recover more slowly than an unconnected population, it is of interest whether such an optimal set point has advantages in the network setting. Moreover, it remains an open question which brain regions ought to have such filter properties.

### Communication channels

Many factors influence the optimal set point: among others, the balance of excitation and inhibition, but also the recurrent connectivity, which is likely to be non-random (Fig. 5b, c). More generally, heterogeneity in the neural and synaptic properties can shape the reaction and the recovery speed individually, and consequently, likely also on a population level. Furthermore, neuromodulation can alter several properties of a network temporarily, or even permanently through synaptic plasticity (Fig. 5c). How heterogeneity and modulation affect the response properties of neural networks remains an important topic for future research.

**Figure 5.**
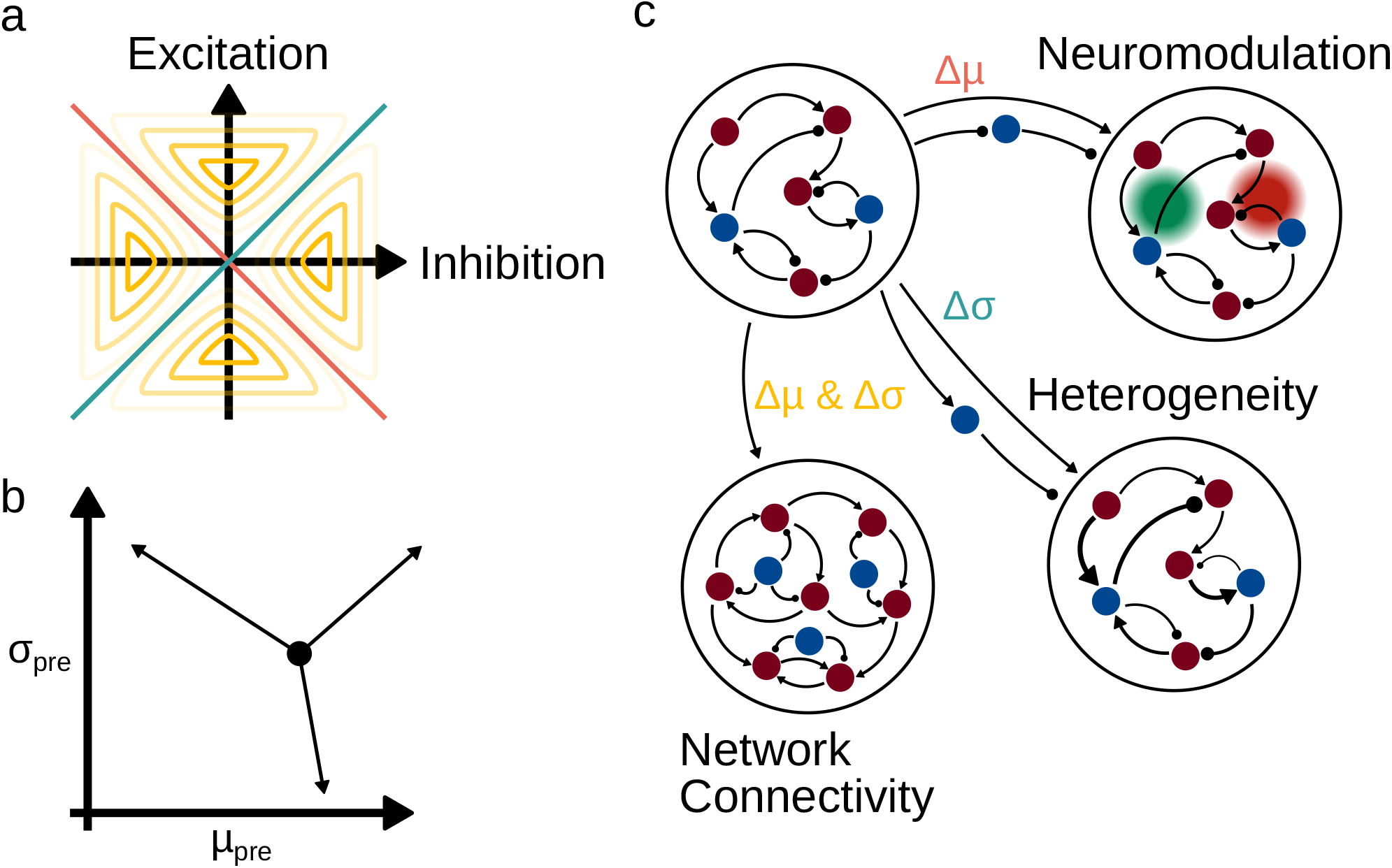
The communication between neural populations is influenced by network characteristics, inter-areal connectivity profiles, and can be modified across timescales. **(a)** The change in excitation and inhibition determines the communication modality. If either or both change, the default modality is Δ*µ &* Δ*σ*. Only if excitation and inhibition increase proportionally (inversely proportional), the communication is through the modality Δ*σ* (Δ*µ*). **(b)** Neuromodulation, synaptic plasticity or other means effect the set point of the network. **(c)** Inter-areal connectivity profiles support different modalities (e.g. Δ*µ*: disinhibition and feed-forward (FF) excitation; Δ*σ*: FF inhibition and FF excitation; Δ*µ &* Δ*σ*: FF excitation). The network characteristics of the receiving network are shaped by its internal connectivity (e.g. convergence or divergence, or the balance of excitation and inhibition), its heterogeneity in neural types. Additionally, these features can be dynamically altered by neuromodulation or synaptic plasticity across various timescales.

While the optimal set point is a property of the receiving network, the communication between networks incorporates more: namely, the modality of communication (Δ*µ*, Δ*σ*, or Δ*µ &* Δ*σ*), features of the activity (persistent or stimulus pulses of varying duration), and the connectivity profile. The characteristics of the response dynamics depend on the features of the communication and can be altered or tuned. One major point is that the response properties were contingent on the modality of the communication. Only changing the mean drive or the fluctuation level is a special case, while most often, both parameters are intertwined (Fig. 5b; Brunel 2000). Here, the projection pattern of the sending population becomes crucial. If projections are only excitatory, both output parameters are affected. On the other hand, the addition of feedforward inhibition of equal strength effectively alters only the fluctuation level. In contrast to feedforward disinhibition, in which solely the mean drive is changed. For the latter two cases, this only holds under the condition of equal strength in the projection strength (Fig. 5a, c). Thus, neural activity could transmit different information based on the projection pattern between networks.

Moreover, the communication among networks may have different features, be it persistent or transient stimuli. With the assumption that a single stimulation pulse encodes one bit of information, the recovery time indicates the information bandwidth of the neural population.

When more than one pulse was presented to the neural population, the recovery sped up in some cases, while in others it slowed down (Suppl. Fig. 2). Those experiments also revealed that neurons exhibit hysteresis (Fig. 3a) that depended on the properties of the pulse trains (number of pulses and pulse duration). We observed that the longest recovery time for transient stimuli depended non-monotonically on the number of pulses as well as their stimulus duration (for a given set point and modality). Therefore, the network’s set point and the communication modality ought to be fine-tuned to the typical stimulus properties of the sending network. Heterogeneity could also play a crucial role in decoding a wider range of stimuli. In contrast to typical stimuli, a surprising event may be encoded predominantly in a fast reaction time, while the recovery speed only plays a subordinate role. These insights dismiss a simplified view of information encoding and make information bandwidth history- and stimulus-dependent (Fig. 3b-d).

### Implications

Long recovery times impose strong limits on how fast neural networks can respond to time-varying signals. The importance of fast reaction time may not be underestimated, as stimuli are constantly reaching the brain. However, this becomes meaningless if the neurons recover sluggishly. This entails the question of which neural mechanisms can calibrate the response properties along both axes - fast recovery while keeping quick reaction properties.

One way out of this problem would be to have a tight balance of excitation and inhibition (Hennequin, Vogels, and Gerstner 2014), but such a balance is untenable in *in vivo* settings. Alternatively, neural networks could rely on compressed sensing to recover the full input despite low sampling frequency (Ganguli and Sompolinsky 2012) using internal models of the input. In the absence of proper internal models, neural networks would have to use the notion of temporal interleaving or spatial interleaving to process fast stimuli. In temporal interleaving, the slow neurons would sample the same signal at different phases, and in spatial interleaving, different phases of the signal are measured in different parts of the brain. The solutions from electronic communication theory are tempting, but it would be necessary to test their biological plausibility.

### Experimental verification and predictions

With the technology currently available to stimulate and record hundreds of neurons simultaneously, it is in principle possible to replicate the simulation experiments *in vivo*. However, heterogeneities in the neural properties or even the connectivity profiles are likely to affect the response dynamics. Moreover, we assumed no short-term synaptic dynamics by using static synaptic properties. The same holds for the neural properties that did not allow for adaptation features. A major challenge is also the comparison of biological neurons and LIF neurons. However, the crucial common ground is the non-linearity due to the reset mechanism. We believe that more complex neuron models or even biological neurons share fundamental properties in the response dynamics, namely that the recovery speed is finite and depends on neuron, network, and even stimulus properties.

Lastly, It should be pointed out that in our simulations, all neurons in a population had the same membrane properties, including the membrane time constant. And yet the neuron populations displayed a large range of reaction and recovery times depending on the experimental setup. The reset mechanism after a spike introduces non-linear effects (Deger et al. 2010), and thus fitting a single or a double exponential function to the response was not applicable (see autocorrelation functions in Suppl. Fig. 3). Although matching the dynamics to a linear second-order filter seemed more promising, certain responses could not be fitted, as the initial response jump exceeded the step input by more than twice. Consequently, we advocate focusing on analyses that are agnostic to a single time scale. Ultimately, the dynamics of neurons, neural populations, and the whole brain evolve on temporal manifolds (Marom 2010). Hence, an interesting line of research would involve temporally more complex stimuli, from coloured noise to experimentally obtained inputs.

## Code availability

The simulation code is available on GitHub (https://github.com/Jiggaboy/responsiveness).

## Author contributions

Hauke O. Wernecke and Arvind Kumar designed the study and interpreted the data; H.W. analyzed the data, developed code, and performed simulations; H.W. wrote the manuscript with contributions from all co-authors.

## 1 Supplementary Figures

**Supplementary Figure 1.**
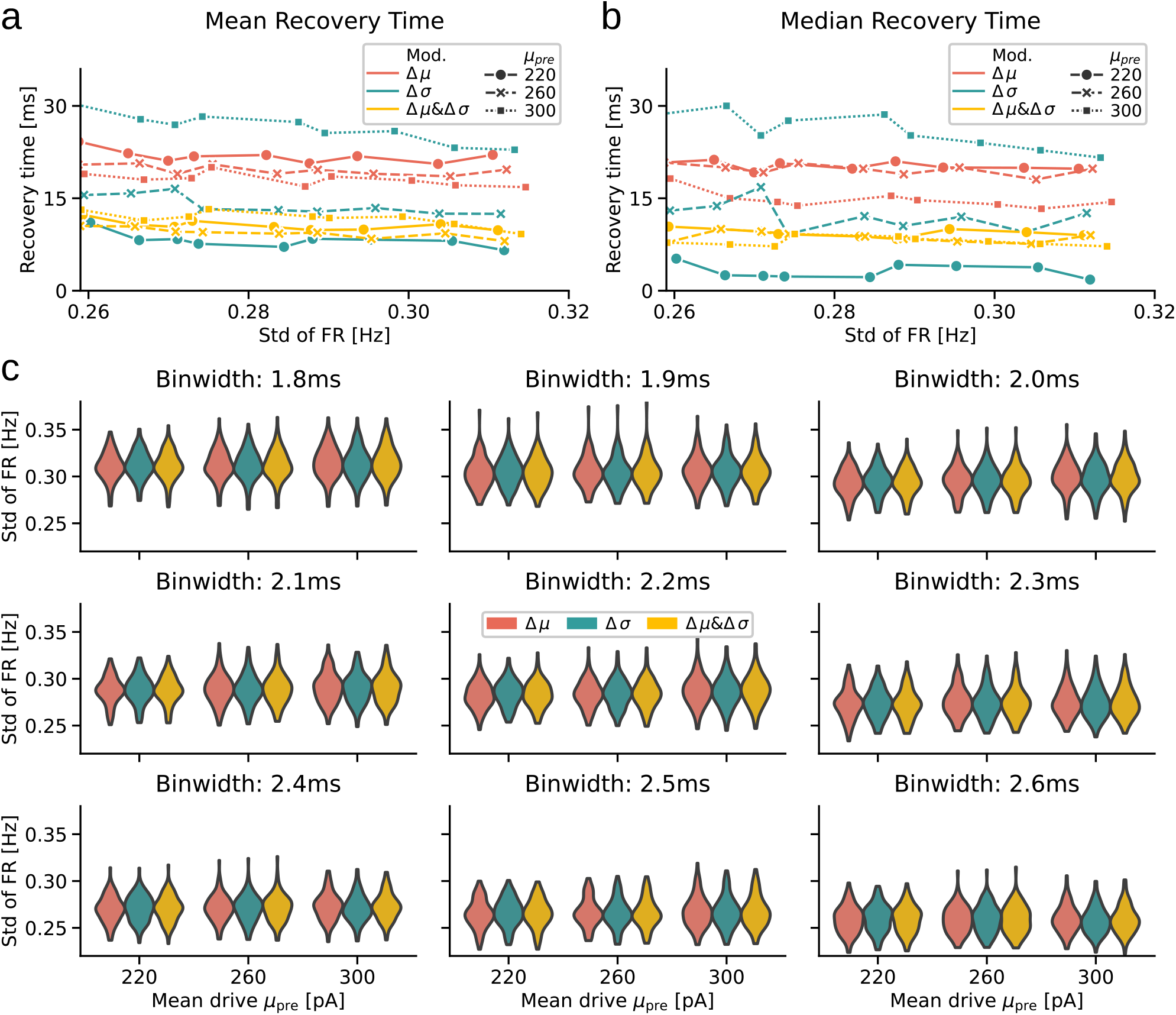
**(a)** Mean recovery time depended on the sample standard deviation Std of the FR (Std quantified after reaching stationary state, averaged across bootstrap samples). **(b)** Same as in (a) for the median recovery time. **(c)** Distribution of the sample standard deviation *Std* across various binwidths, modalities and set points.

**Supplementary Figure 2.**
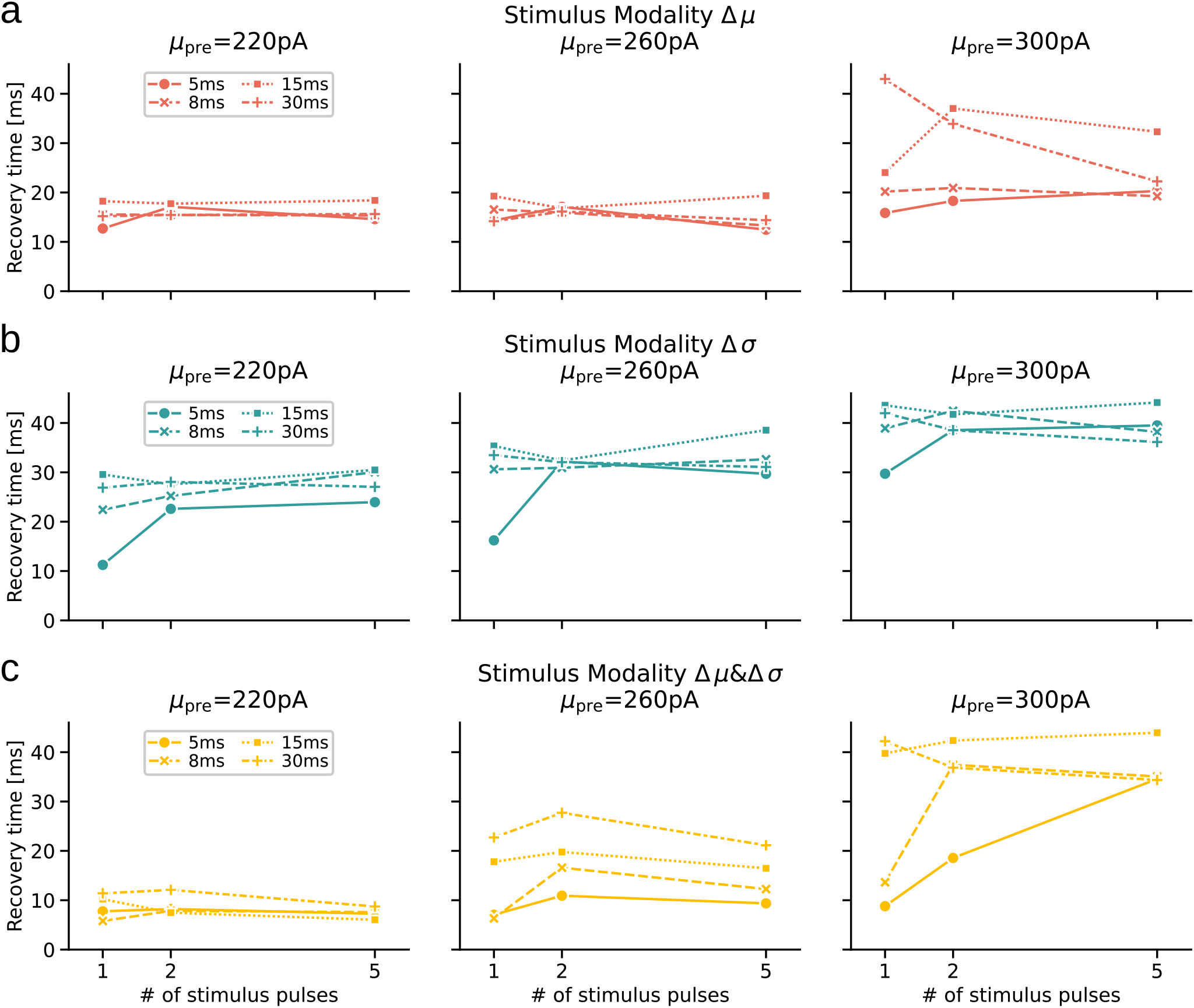
Stimulus duration affected the recovery time across modalities. **(a)** Input modality Δ*µ*: The recovery time depends non-monotonically on the number of stimuli. This dependence revolved around the set point as well as the stimulus modality. Three set points are shown: *µ*_pre_ = 220pA (left), *µ*_pre_ = 260pA (middle), *µ*_pre_ = 300pA (right). **(b)** Same as in (a) for stimulus modality Δ*σ*. **(c)** Same as in (a) for stimulus modality Δ*µ &* Δ*σ*.

**Supplementary Figure 3.**
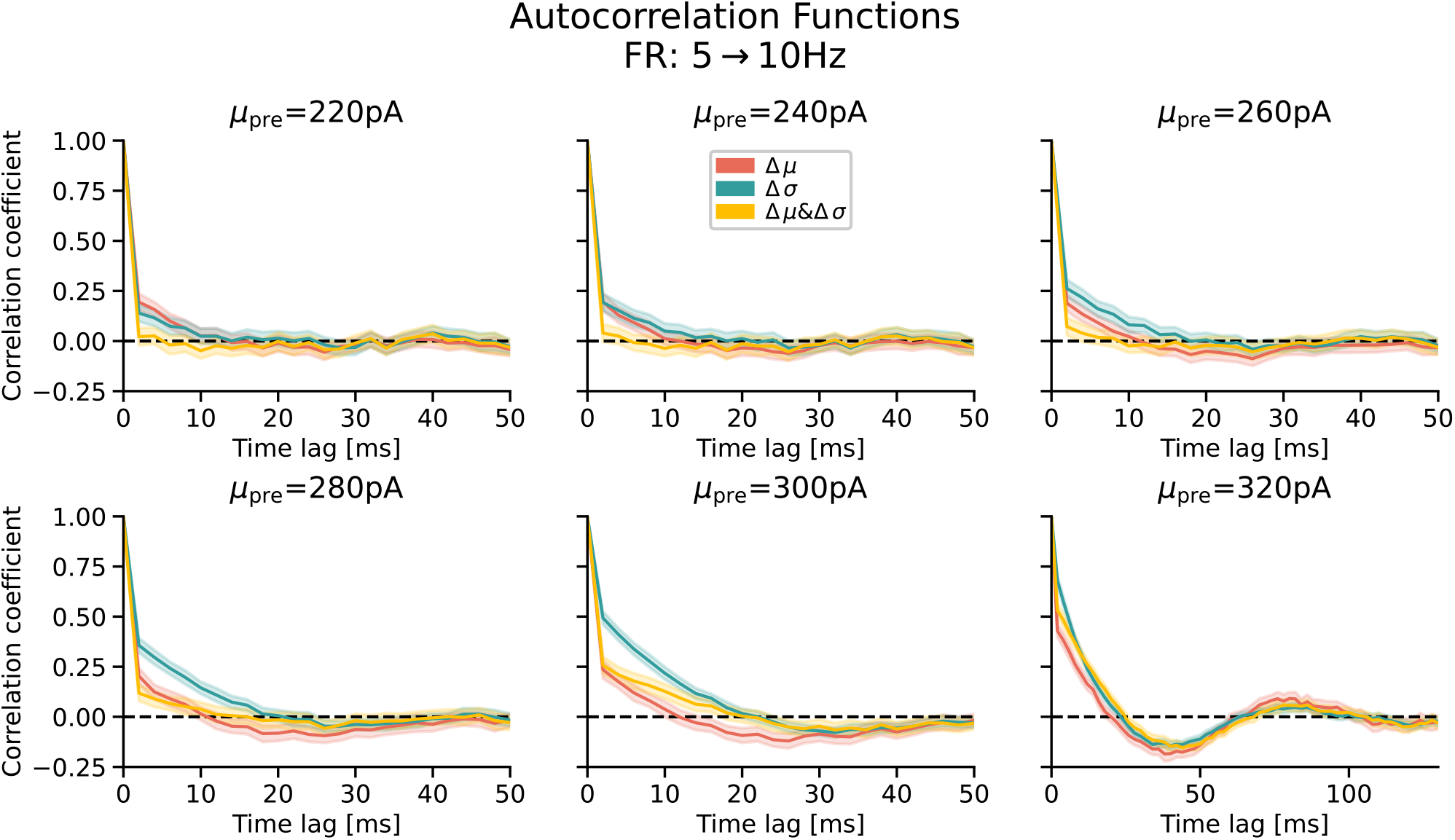
Autocorrelation functions of the neural activity across modalities and set points.

**Supplementary Figure 4.**
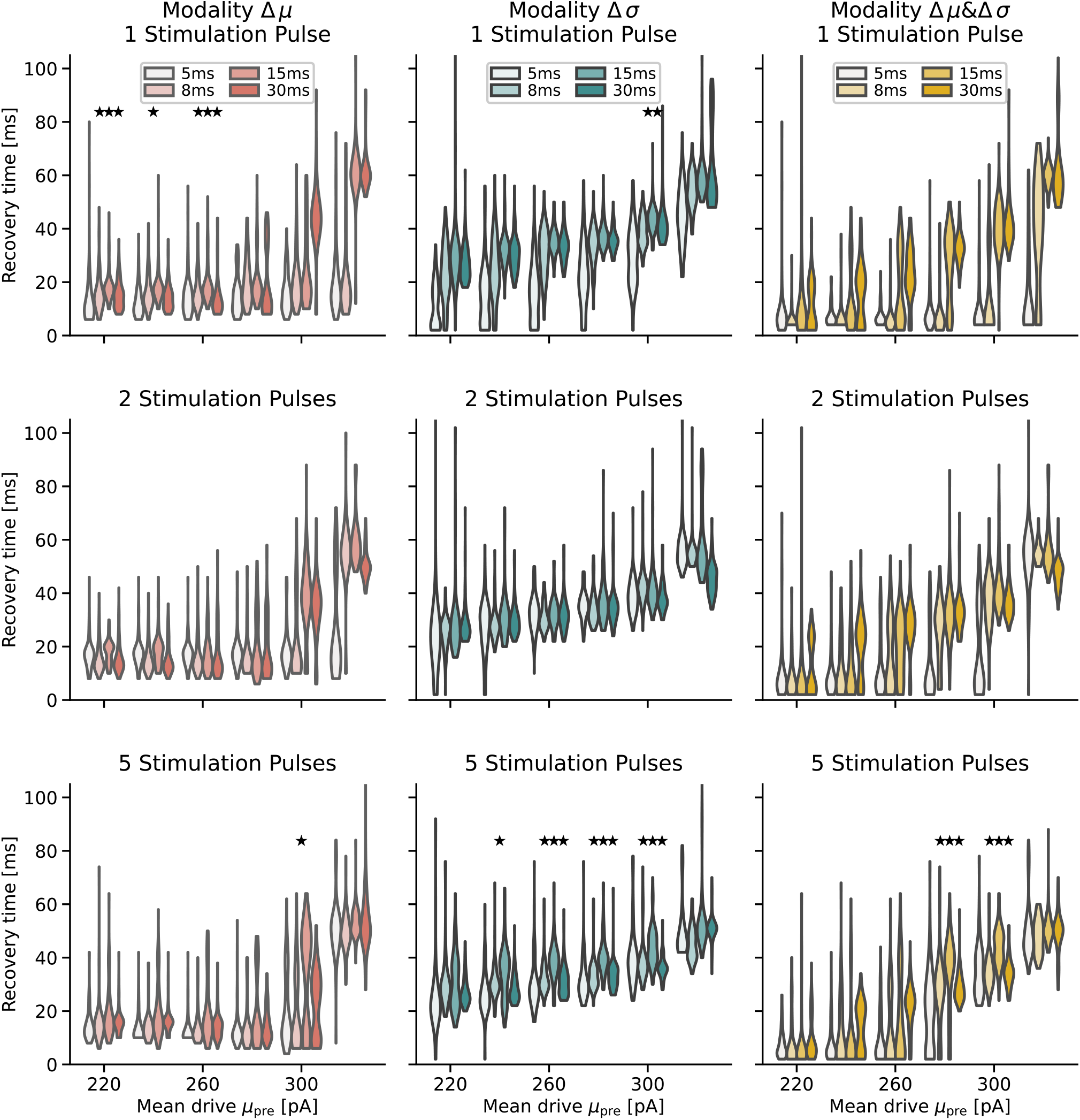
Distribution of recovery times across set points, stimulus duration and number of stimulus pulses (rows) across stimulus modalities (columns). Asteriks indicate level of significance that the stimulus duration of t_stim_ = 15ms is slowest to recover (for details see Methods). *p* values (from left to right): one stimulation pulse: 1.07*e* −06, 6.72*e* −05, 4.652*e* −06 (Δ*µ*); 4.44*e*−05 (Δ*σ*); five stimulation pulses: 7.36*e*−05 (Δ*µ*); 8.57*e*−05, 8.17*e*−13, 4.34*e*−18, 2.92*e* − 08 (Δ*σ*); 2.73*e* − 09, 6.83*e* − 14 (Δ*µ &* Δ*σ*);

